# The patterns of codon usage between chordates and arthropods are different but co-evolving with mutational biases

**DOI:** 10.1101/2023.03.30.534958

**Authors:** Ioanna Kotari, Carolin Kosiol, Rui Borges

## Abstract

Different frequencies amongst codons that encode the same amino acid (i.e. synonymous codons) have been observed in multiple species. Studies focused on uncovering the forces that drive such codon usage showed that a combined effect of mutational biases and translational selection works to produce different frequencies of synonymous codons. However, only few have been able to measure and distinguish between these forces that may leave similar traces on the coding regions. Here, we have developed a codon model that allows the disentangling of mutation, selection on amino acids and synonymous codons, and GC–biased gene conversion (gBGC) which we employed on an extensive dataset of 415 chordates and 191 arthropods. We found that chordates need 15 more synonymous codon categories than arthropods to explain the empirical codon frequencies, which suggests that the extent of codon usage can vary greatly between animal phyla. Moreover, methylation at CpG sites seems to partially explain these patterns of codon usage in chordates but not in arthropods. Despite the differences between the two phyla, our findings demonstrate that in both, GC–rich codons are disfavoured when mutations are GC–biased, and the opposite is true when mutations are AT–biased. This indicates that selection on the genomic coding regions might act primarily to stabilise its GC/AT content on a genome–wide level. Our study shows that the degree of synonymous codon usage varies considerably among animals, but is likely governed by a common underlying dynamic.

**Significance statement:** The reasons for the differential usage of codons encoding for the same amino acid has puzzled scientist for decades. By examining the frequencies of synonymous codons in different species, this study presents a novel model that sheds light on the underlying factors that drive differences in codon usage between chordates and arthropods. Our analysis unveiled more extensive codon usage patterns in chordates compared to arthropods. Despite differences between the phyla, the study highlights that genome– wide selection acts to balance mutational biases, as GC-rich codons are less favoured under GC-biased mutations, while the opposite holds true for AT-biased mutations. This research provides valuable insights into our understanding of the complex interplay between mutational biases and selection forces in shaping the variation at the synonymous sites, and has important implications for future studies of genome evolution and adaptation.

## 1 Introduction

Synonymous codons encode for the same amino acid and are expected to be neutral and used interchangeably in the genome. However, synonymous codons appear at different frequencies across protein–coding genes. This preferential usage of synonymous codons is called codon usage bias and is widely spread across the tree of life (Subramanian, 2008). In order to explain the differential use of synonymous codons, two main narratives have been proposed: mutational biases and translational selection (Duret, 2002; Hershberg and Petrov, 2008). Translational selection is expected to affect highly expressed genes and correlates well with tRNA abundance to increase translation efficiency, and has been found to bias codon usage in a number of taxa, e.g., *Drosophila* (Shields et al., 1988; Akashi, 1994; Bierne and Eyre-Walker, 2006), *C. elegans* (Duret and Mouchiroud, 1999), as well as in *E. coli* (Ikemura, 1981). On the other hand, mutational biases have been found to shape codon usage bias according to the base composition of the whole genome and GC–content in the third codon position (e.g., yeast (Sharp et al., 1995), and vertebrates (Urrutia and Hurst, 2001; Rao et al., 2011) including humans (Sueoka and Kawanishi, 2000)).

The exact causes of codon usage bias are still widely debated, but it is of common agreement that it arises from the interplay of selection, mutation, and genetic drift (Bulmer, 1991). Studies have found that mutational forces are acting alongside the well–established effect of translational selection on codon usage bias in *Drosophila* (Kliman and Hey, 1994). Similarly, a correlation between preferred codons and tRNA levels was discovered in several vertebrate species, showing that weak translational selection operates alongside mutations (Doherty and McInerney, 2013). Additionally, Galtier et al. (2018) recently described how GC–biased gene conversion (gBGC) – a meiotic recombination–associated bias that favours GC– over AT–alleles (Marais, 2003) – has a widespread effect on codon preferences across animal species. Therefore, the debate has changed from identifying the forces responsible for codon usage bias to determining which of these forces are more significant.

The assessment of codon usage bias has traditionally relied on heuristic methods, e.g., codon usage indices. The most commonly used indices include the Codon Adaptation Index (CAI) (Sharp and Li, 1987), the Relative Synonymous Codon Usage (RSCU) values (Sharp and Li, 1986) and the GC–content at the third position of synonymous codons (e.g., Galtier et al. (2018)). These cluster the different codons in a small number of categories, usually as preferred and non-preferred, but fail to fully account for the effects of mutation and selection processes. To disentangle these effects, mechanistic approaches that rely on population genetics models have been proposed. A maximum likelihood approach includes the multi-allele model (Zeng, 2010), which attempts to categorise codons into four classes, and two Bayesian approaches, FMutSel (Yang and Nielsen, 2008) and ROC SEMPPR (Gilchrist et al., 2015), which quantify selection and mutation’s impact on codon preference. However, none of these approaches incorporates gBGC as a factor in codon usage bias.

Previous research was restricted to a few model organisms in animals and mainly focused on highly expressed genes. Most extensive interspecific studies of codon usage are focused on yeast (LaBella et al., 2019) and bacteria (Sharp et al., 2005). With advancements in sequencing technology, whole genomic coding sequences from a wider range of species are now accessible. This led recent studies to shift their focus on inter-species variation in animals as well. Doherty and McInerney (2013) analysed a range of vertebrate species, while Galtier et al. (2018) expanded beyond vertebrates and included species across the whole animal kingdom. However, data now exists that allows a more detailed comparison of the patterns of codon usage across phyla.

In this study, we devised a mechanistic model based on the mutation-selection Moran model (Moran, 1958), which explains the fixed differences between species based on population genetics forces. To estimate these forces, we developed a Bayesian estimator called DECUB (Disentangling the Effect of Codon Usage Bias) which quantifies the joint effect of mutations, selection and gBGC across the whole genome. We focused on chordates and arthropods as these two phyla are well studied with sufficient genomic sequence availability, have both diverged during the Ediacaran Period (635-538 MYA) (Dos Reis et al., 2015) and the main driver of codon usage bias has been attributed to translational selection in arthropods but mutational biases in chordates. We employed our model on coding sequences from over 600 species belonging to these phyla to (i) disentangle and evaluate the effects of the aforementioned confounding forces of codon usage, and (ii) compare their patterns between the phyla. We found that codon usage bias is more extensive in chordates compared to arthropods, and that genome-wide codon usage in both taxa is co-evolving with mutational biases.

## 2 Results

### 2.1 Modelling the evolution of codon frequencies

In this study, we created a Moran model with mutations, GC–bias, and selection to model the codon frequencies along the genome. Mutations are modelled similarly to the general time–reversible (GTR) substitution model (Tavaré, 1986) and GC–bias is incorporated to capture the effects of GC–biased gene conversion (gBGC). The joint effects of these on a given codon *I* = *i*_1_*i*_2_*i*_3_ are summarised in the mutational coefficient of all three nucleotides *β_I_* = *β_i_*_1_ *β_i_*_2_ *β_i_*_3_. Selection acting on said codon is modelled as a relative fitness coefficient *Φ_I_*.

As we are interested in capturing the variation at the inter–species level, we used the stationary frequencies of the fixed sites (supplementary text S1, S2), which can be described for each codon, as

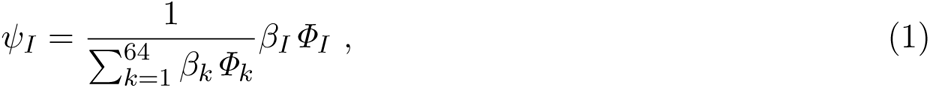

where the denominator is set such that the stationary frequencies add up to 1.

Using the stationary distribution in eq. (1), we developed a software called DECUB (Disentangling the Effects of Codon Usage Bias), which takes as input codon counts from a given taxon (e.g., population, species) and infers the mutational coefficient of each nucleotide and the fitness coefficients of each codon using a Bayesian framework. All mutational coefficients are normalised based on adenine (A) since the effect of gBGC does not influence mutational biases towards A. Similarly, as methionine is an essential amino acid encoded by a single codon and is therefore not confounded with codon usage bias, all fitness coefficients are normalised based on the codon ATG. The general model presented in (1) can potentially assume a fitness coefficient per codon. However, after performing extensive simulations, we found that the model is unidentifiable after 53 fitness coefficients (supplementary fig. S1).

### 2.2 Assessing the evolutionary significance of the model estimates

It is known that variation in the strength of gBGC and mutations vary greatly in the genome, and spatially heterogeneous selection on codon usage bias scales with expression (e.g. Gilchrist et al. (2015); Cope and Shah (2022)). To establish that our model accurately quantifies the strength of these underlying processes, we conducted simulations using genome-wide data from humans (*Homo sapiens*) and fruit flies (*Drosophila melanogaster*) as case studies. We generated codon counts by incorporating variation in mutations, gBGC, and codon fitnesses per gene, intending to more realistically reflect observed variation across coding sequences (CDS) in both species. While our model cannot capture this heterogeneity or assess its position, we found that DECUB measures the average value of those forces across the collection of genome-wide codon data.

Our analyses revealed highly significant Spearman’s *ρ* rank correlations between the mean true fitness coefficient across all genes and the estimated values (supplementary fig. S2, S3), indicating that our estimates capture biological signal consistent with gene-wide averages. Furthermore, relative errors below 20% indicate strong concordance between simulated and estimated values (supp. fig. S4). Regarding the mutational biases, in cases of more homogeneous GC-biased gene conversion (gBGC), like in *Drosophila*, our estimations correlated significantly with the mean of the simulated values (supp. fig S2). However, in the presence of recombination hotspots, as is the case in humans, where areas of the genome experience extreme values of gBGC, these correlations are lower (supp. fig. S3). These hotspots are typically found in only 1-2% of the genome (Glémin et al., 2015), predominantly outside coding regions (Myers et al., 2005). However, in our simulations, we assumed 2% of hotspots solely in coding regions, resulting in a much higher level of heterogeneity than observed in reality. Consequently, a majority of these extreme values were concentrated in a small portion of the dataset. Due to this disparity, we find that the estimated mutational biases for GC alleles (*β_C_* and *β_G_*) tended to be closer to the median of the distribution of simulated values rather than the mean (fig. S5). However, this did not bias the estimation of any GC-rich codon fitness coefficients, proving the capability of our model not only to capture mutational biases and fitness coefficients but also to disentangle them based on pooled codon counts, accurately measuring their mean value across coding genes.

### 2.3 Chordates have more pronounced patterns of codon usage than Arthropods

To characterise the patterns of codon usage in arthropods and chordates, we used codon counts from genome-wide coding sequences of 606 species: 415 chordates and 191 arthropods. All species are encoded as per the standard genetic code. We set up our inferences by using an amino acid mapping, which sets a fitness coefficient per amino acid, plus one per stop codon (total 23 *Φ* categories). Figure 1 shows the estimation of each mutational and fitness coefficient on a logarithmic scale. Although the estimates were distributed similarly, chordates were more homogeneous in their estimates across the species studied compared to arthropods.

**Figure 1:**
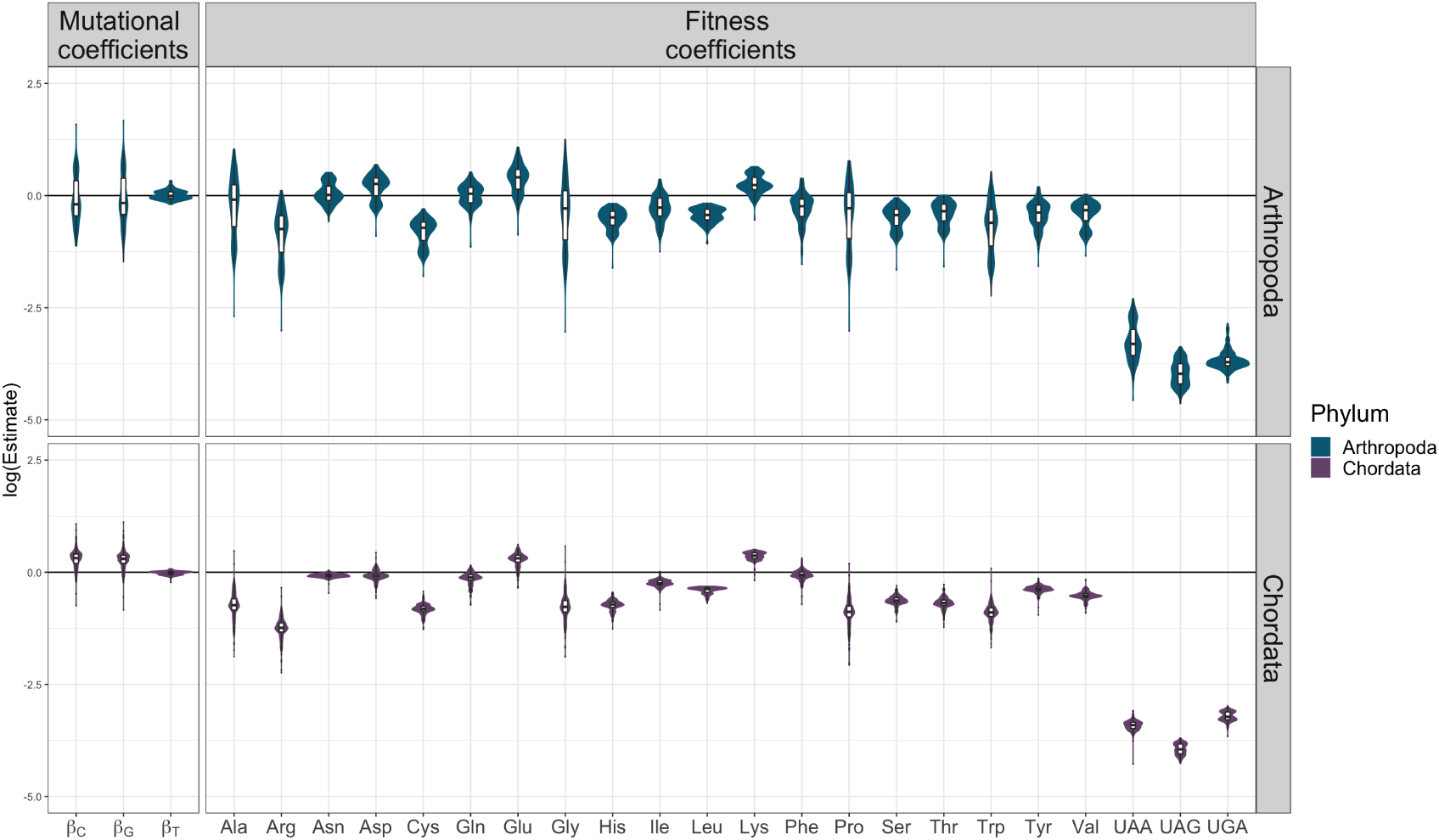
Estimation of mutational and fitness coefficients under an amino acid mapping. Logarithmic scale of each estimate which shows positive values if a coefficient is estimated higher than the reference one or negative if lower. The mutational coefficients were normalised with *β_A_* which does not include the effects of GC-biased gene conversion (gBGC) and the fitness coefficients were normalised by the single codon for methionine.

The estimated mutational biases of thymine (*β_T_*) are close to 0, therefore almost equal to *β_A_*, the mutational effects of which were used to normalise all mutational coefficients. Moreover, as would be expected, we see greater variation in the mutation coefficients of GC alleles, which are driven by the combined effect of mutations and gBGC. The estimation of the fitness coefficients highlights the differential usage of each amino acid, and because we did not confine our analyses to the sense codons, it informs us on how deleterious the stop codons are in comparison to sense codons. Arthropods show a preference for TAA as their stop codon, despite its median fitness being 13 times lower than that of the lowest amino acid (arginine). On the other hand, chordates prefer TGA as their stop codon, with a median that is 7.3 times lower than that of the lowest amino acid (also arginine).

The observation that the coefficients only change slightly between phyla reflects how fitness coefficients inherently mirror the structure of the genetic code, representing the usage of each amino acid relative to methionine along the coding regions. We use the term “fitness” to convey this relative preference and in reference to its function in the Moran model. Note that the fitness coefficients can contain forces beyond selection, such as other mutational and recombination biases that have not been directly modelled. However, we can use these estimates as a baseline for our subsequent analysis. By building upon these estimates, we can introduce additional fitness coefficients to investigate genome-wide patterns that extend beyond the genetic code’s structure (i.e., codon usage bias) and the mutational biases we have modelled.

The amino acid mapping expresses the genetic code and is a natural approach to modelling fitness effects; however, it ignores variation between synonymous codons, thus disregarding codon usage bias. To identify codons needing their own fitness coefficient, we used an approach based on the posterior predictive checks by Gelman and Hill (2006), where we compared the error between the empirical codon frequencies and the predicted ones from our model. Figure 2 shows the percentage of species whose estimates deviate from each empirical codon count. In arthropods, most of the observed variation can be sufficiently explained with the amino acid mapping. However, more codon categories are clearly needed to account for the more extensive variation amongst the synonymous codons in chordates. It is important to note that these added fitness coefficients aim to capture codon-specific effects, which are most likely due to codon usage bias, but can also encompass some amino acid effects which we cannot disentangle. For this reason, we have henceforth used the term “codon usage bias” only when we explicitly mention this phenomenon, and employ “codon usage” or “codon preferences” when discussing codon-specific fitness estimates that encompass codon usage bias.

**Figure 2:**
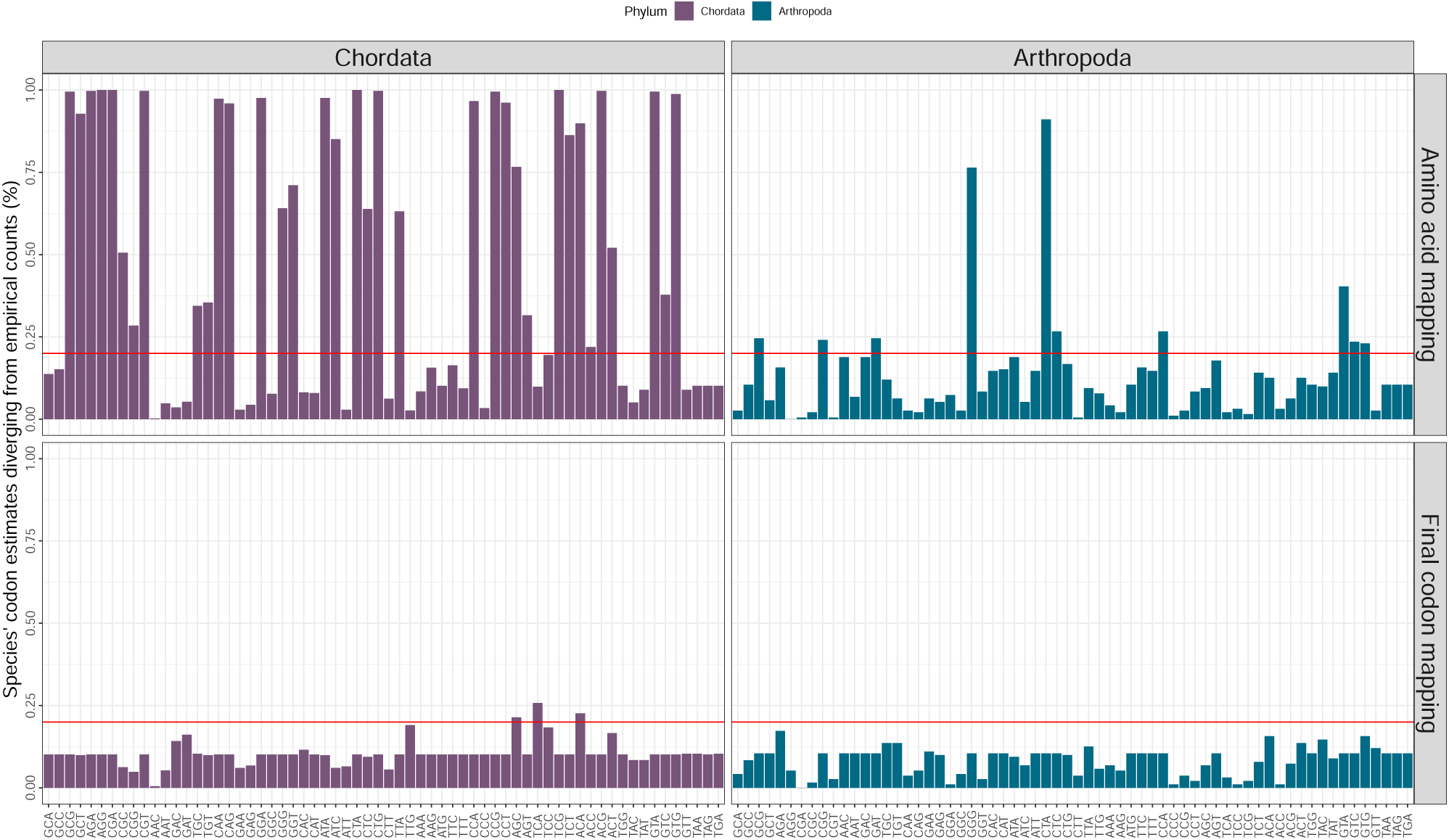
Error plots between predicted and empirical codon frequencies in chordates and arthropods, differences between the amino acid mapping and the final codon mapping. Assuming no variation amongst synonymous codons (amino acid mapping, i.e. one *Φ* per amino acid), showed a more extensive codon usage pattern in chordates compared to arthropods. In chordates (left, purple) 35 codons out of 64 had more than 20% of their species being predicted outside the empirical distribution, whereas arthropods (right, blue) had overall smaller errors and only 10 codons exceeding the 20% threshold. We proceeded to minimise these errors per codon by adding more fitness coefficients. Each new mapping adds a new fitness coefficient per amino acid to the codon that had the highest error in the previous mapping. The final codon mapping (shown in the bottom plot) adds 29 extra coefficients to the chordates and 14 to arthropods, resulting in a total of 53 and 37 coefficients, respectively. We note that 3 codons still exceed the 20% threshold, as the model reached the identifiability threshold in chordates.

We proceeded to add more fitness coefficients in a step-wise manner to the codons whose estimates do not fit that of the empirical distribution in more than 20% of species. First, we added a fitness coefficient to the codon of each amino acid that had the highest percentage of error. We proceeded to repeat this procedure until all variation was below the 20% threshold, resulting in a final phylum-specific mapping (fig. 2). We note that chordates reached the identifiability threshold with 3 codons exceeding 20%; however, the errors of these were relatively low, not exceeding 25%. We employed the Bayesian and Deviance Information Criteria (BIC and DIC) (Schwarz, 1978; Spiegelhalter et al., 2002), where we aimed to find the best fitting mapping. In all species but seven arthropods, the last codon mapping was the optimal one (supplementary table S2). Hence, the final chordate mapping has 52 fitness coefficients compared to arthropods that are modelled with 37.

Although chordates exhibit greater variation in synonymous codons, this pattern is comparatively more uniform across species compared to species within arthropods, which display less extensive bias but greater heterogeneity within their taxa (supp. fig. S6). Between arthropods and chordates, however, the final mappings show clear differences in patterns of codon usage, with chordates needing over twice the number of extra coefficients compared to arthropods (29 vs. 14 extra codon fitness coefficients).

### 2.4 Fitness coefficients balance mutational effects

We established that chordates and arthropods show different extent of codon usage; additionally, the mutation and fitness coefficients vary considerably within these phyla (fig. 1, S6). To understand these differences, we focused our analyses on the main representative chordate classes (mammals, birds, reptiles and amphibians, and fish), which comprise 97% of the dataset. Similarly for arthropods, the subsequent analyses focus on Diptera, Lepidoptera, and Hymenoptera (flies and mosquitoes, butterflies and moths, and ants, wasps and bees, respectively), which make up for 74% of the arthropod dataset, and more specifically more than 83% of the collected insect species (fig. 6, table S3).

**Figure 3:**
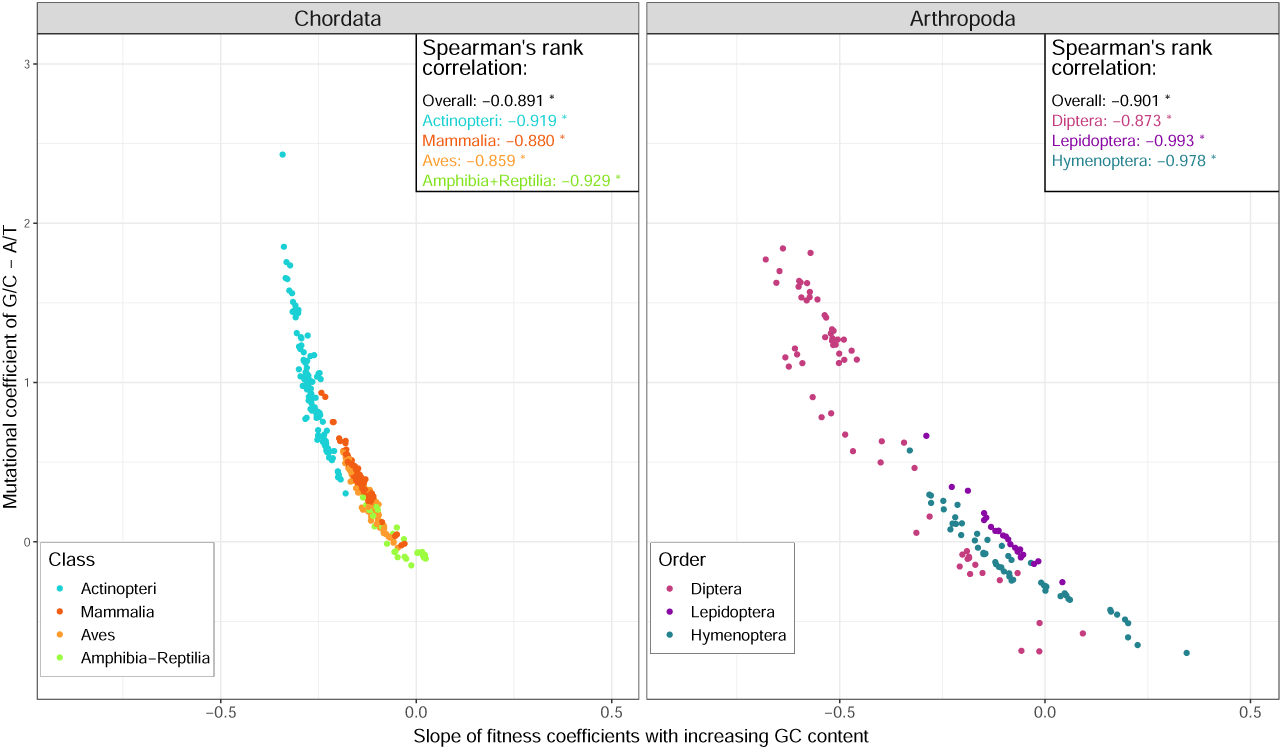
Negative correlation between mutational biases towards GC and increasing GC– content of codon fitness coefficients in chordates and arthropods. The larger the difference between the mutational coefficients of GC and AT nucleotides, the more negative the slope of fitnesses is, i.e. the less favourable a codon will be if it is richer in GC–content. In both chordates (left) and arthropods (right), this relationship is inverse with significant Spearman’s rank correlation coefficients *ρ*, both overall and per inner taxa (box in the upper right corner of each graph). The colours represent the different taxa plotted, the classes fish (blue), mammals (red), birds (orange), and amphibians & reptiles (green) in chordates and the orders Diptera (flies & mosquitoes, pink), Lepidoptera (butterflies & moths, purple), and Hymenoptera (ants, bees & wasps, turquoise) in arthropods.

**Figure 4:**
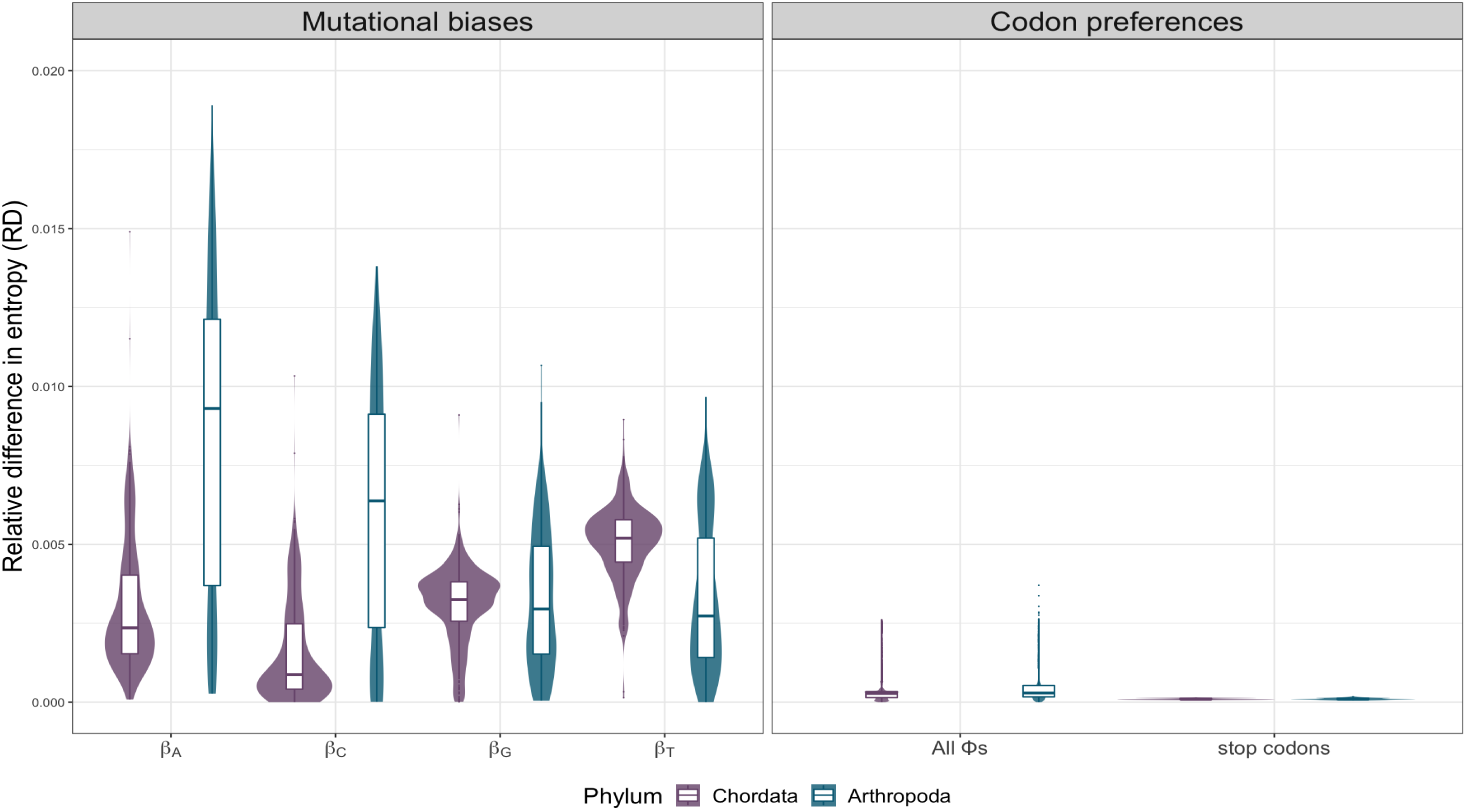
Relative difference in entropy measuring the evolutionary effect of each coefficient in arthropods and chordates. Violin plots with boxplots of the relative difference in entropy of every species, both in chordates (purple) and arthropods (blue). The entropy difference for the mutational biases (left) measures the effect a perturbation of 10% of each *β* has on the stationary distribution. For the fitness coefficients, the summarised effects of all sense and all stop codon fitness coefficients can be seen on the right. The effect of mutational effects are different between chordates and arthropods but both are multiple orders of magnitude higher than codon fitnesses (18.6*×* and 31.7*×* higher in chordates and arthropods, respectively)

**Figure 5:**
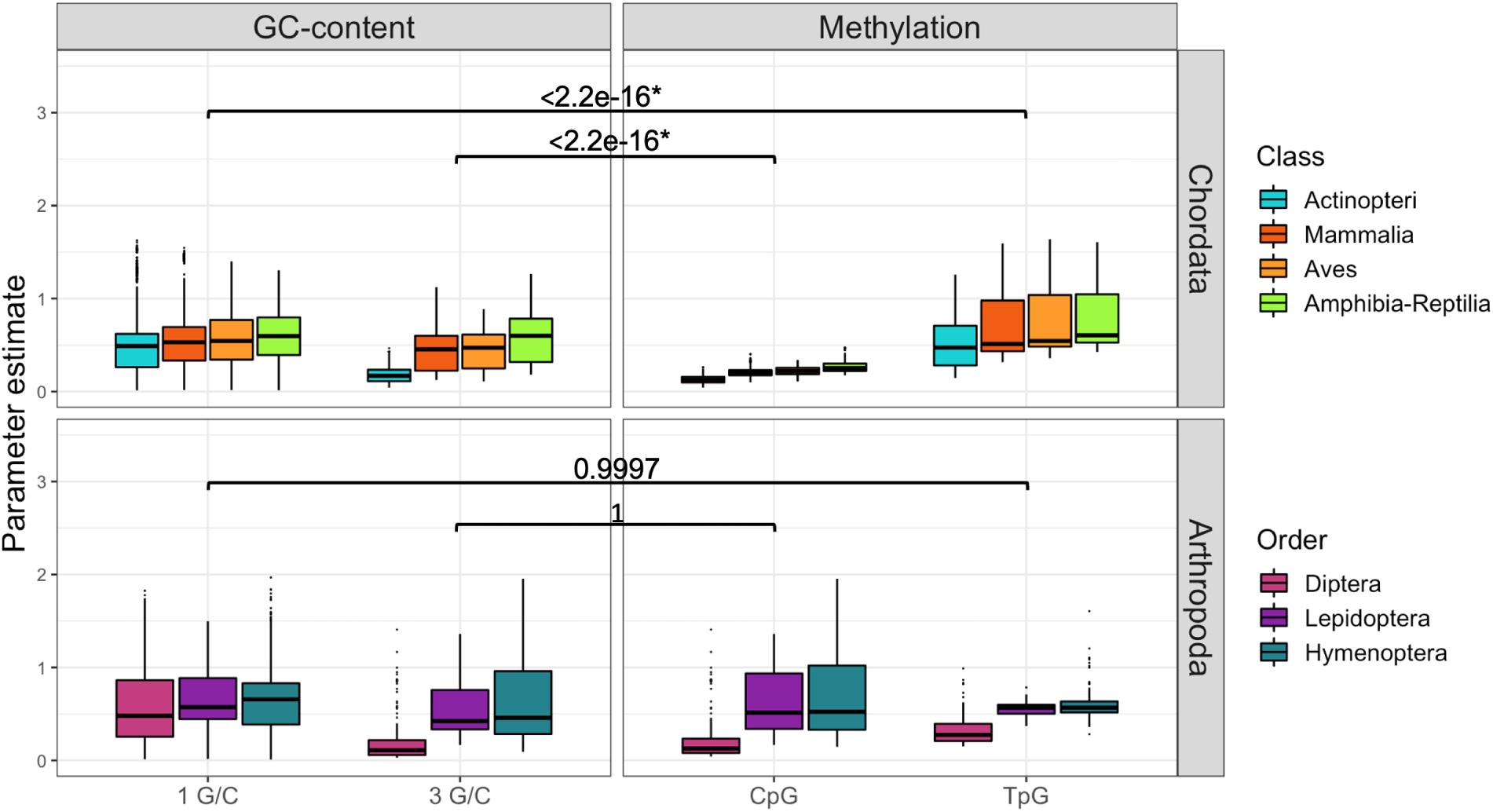
Differences in the fitness of codons with CpG sites compared to their expected GC content. CpG deficit and TpG surplus are expected due to methylation. The non-parametric Wilcoxon signed-rank test was used to compare CpG/TpG sites with their expected estimate due to GC content in chordates and arthropods. Significant results are shown with an, colours represent the different taxa.

**Figure 6:**
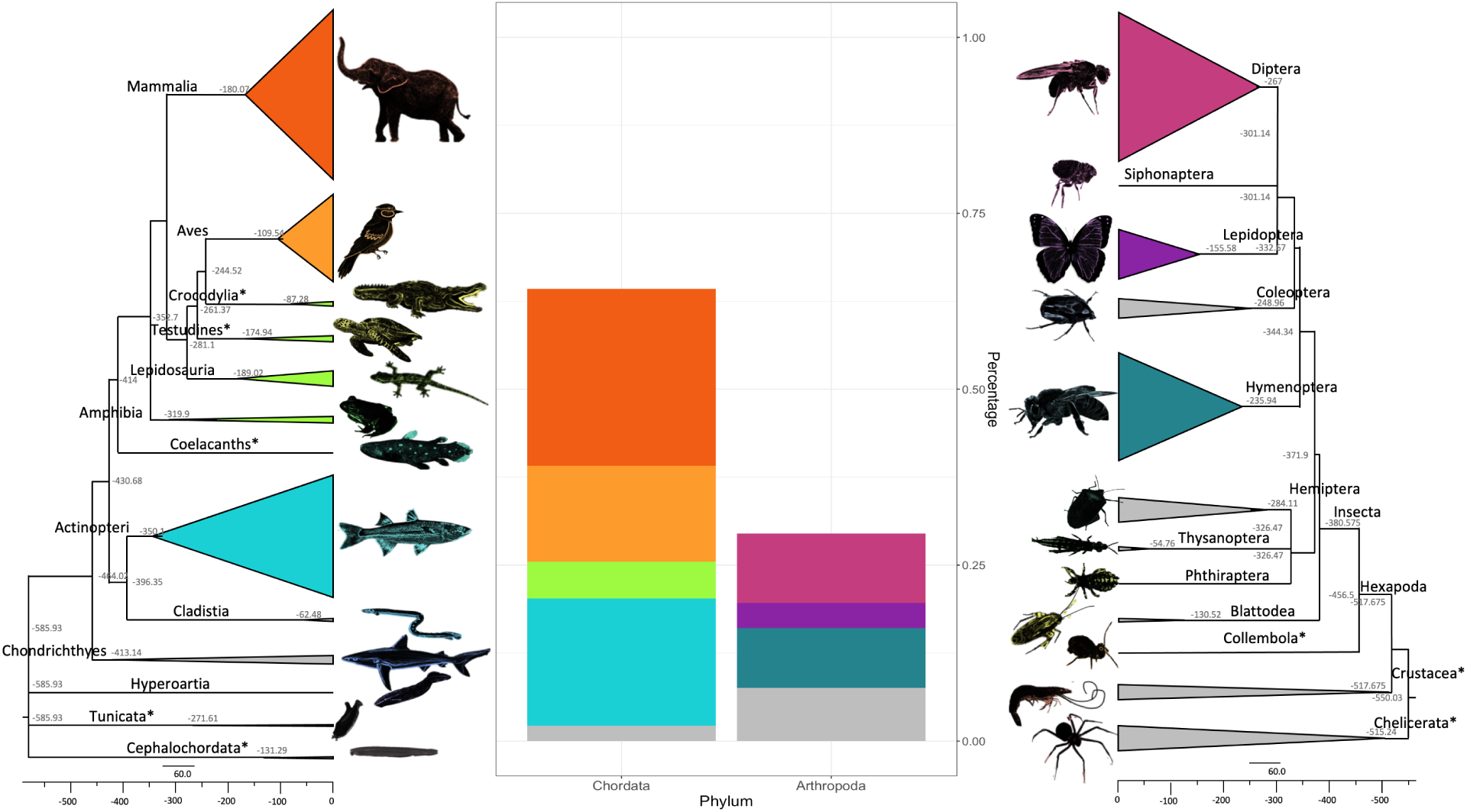
Phylogenetic reconstruction and relative counts of species in the dataset, belonging to Chordata and Arthropoda. The phylogenetic tree on the right represents the 415 chordates species spanning over 13 taxa, while the one on the left represents the arthropods, which summarises 191 species belonging to 12 taxa. For chordates, the taxa with an asterisk next to their names represent a class that was named conventionally by a more commonly known name (e.g. giving the name of the subphyla Tunicata and Cephalochordata to the correspondent class name), or the specific Class has not an official name yet on NCBI (Schoch et al., 2020), therefore was named by its most represented Order (e.g. Crocodylia, Testudines, Coelacanths). In arthropods, we recovered 10 main orders belonging to hexapods (note Collembola* is named here after the most recognised class name), and also summarised the orders found in the subphyla Chelicerata (spiders and scorpions) and Crustacea (crustaceans) which are shown on the phylogeny as a group named after their subphylum followed by an asterisk. The evolutionary time–scales were recovered using TimeTree (Kumar et al., 2022) and visualised with FigTree (Rambaut, 2018), with both trees sharing similar timescales (approximately diverged around 550-600 million years ago, congruent with Dos Reis et al. (2015)). Finally, the middle graph shows the proportions of the selected studied taxa.

In our model, mutational biases also express gBGC. We investigated its role in codon usage by correlating the GC–content of each codon with the bias of mutations towards GC alleles. Figure 3 shows the slope of the codon fitnesses with increasing GC–content for each codon against the difference between GC– and AT– mutational effects for all taxa studied in chordates and arthropods. Both chordates and arthropods have significant negative correlations (Spearman’s *ρ* rank correlation of *−*0.891 and *−*0.901 respectively; *p*–value *<* 2.2 *×* 10*^−^*^16^). These coefficients were obtained after correcting for phylogenetic non-independence using phylogenetic contrasts (Felsenstein, 1985). These negative correlations indicate that the larger the mutational bias towards GC (compared to AT), the less favourable (i.e. the more negative the correlation coefficient) a codon is as its GC–content increases (supp. fig. S7, S8). Additionally, these patterns do not seem to be an artefact of the model, as upon integrating new coefficients, the newly added codons do not show an association with their nucleotide composition (supp. fig. S9), and both GC–only and AT–only codons were chosen in both phyla.

A negative slope between the mutation bias towards GC and the preference for GC–rich codons was also observed among chordate and arthropod groups (all *p*–values *<* 8.4 *×* 10*^−^*^6^). Fishes have the steepest negative slope (*> −*0.25), and along with reptiles, they show the highest correlation coefficients (*−*0.929 and *−*0.919, compared to *−*0.880 and *−*0.859 in mammals and birds, respectively), although all taxa are correlated to a similar degree. Finally, reptiles and amphibians have more homogeneous AT vs. GC mutation preferences, showing a slope very close to 0. Dipterans show some of the largest variation between the mutational differences (*−*1 to 2) with codon fitness slopes ranging from around 0 to *−*0.75. We observed in hymenopterans that AT mutations were equal or sometimes higher than GC, which is reflected in their fitnesses showing a rise (slope that reaches approximately 0.3) in their estimates as GC content increases. Finally, all arthropod taxa were negatively correlated to a similar degree (*−*0.993, *−*0.978 and *−*0.873 in Hymenoptera, Lepidoptera and Diptera, respectively).

To gain further insight into the relative importance of mutational biases and fitness coefficients in determining the patterns of codon usage in chordates and arthropods, we performed a sensitivity analysis. We used the Shannon entropy (Shannon, 1948), as this measure has been previously used to study patterns of codon usage (Suzuki et al., 2004) and is effective in assessing the likelihood of the codon frequencies changing due to variations in the population parameters. We varied the estimated mutation and fitness coefficients by *±*10% and captured the difference in entropy caused to the stationary distribution of the model (see Methods for more details). Figure 4 shows the differences in entropy of each *β* and *Φ* coefficient. It is evident that the mutational biases had a much larger effect on the entropy, with *β_T_* having the largest effect in chordates, as 10% variability results in 0.51% change on the stationary distribution. This effect is 18.6 times larger than the effect of codon fitnesses with an average change of 0.027% (excluding the stop codons). In arthropods, this effect is even larger, with *β_A_* affecting 31.7 times more compared to the effect of the fitness coefficients (0.9% vs 0.029%).

### 2.5 CpG/TpG sites in chordates affect codon usage patterns

CpG/TpG sites are known to affect codon usage patterns in humans (Scaiewicz et al., 2006). As we observed a strong relationship between the codon GC–content and their fitnesses, we aimed to test how much of it could be explained by CpG/TpG sites in chordates and arthropods. Methylated CpG dinucleotides are more likely to change into TpG dinucleotides compared to hypomethylated ones, therefore resulting in a depletion of CpG sites and a surplus of TpG (Simmen, 2008; Miyahara et al., 2015). We employed a one-sample Wilcoxon signed-rank test to test whether CpG (and TpG) codons are less (more) favourable than what is expected due to their GC–content. In chordates, we observed significant differences between the dinucleotides’ coefficients and their expected GC content (*p*–value *<* 2.2*×*10*^−^*^16^), while arthropods show insignificant differences (fig. 5). After correcting for multiple testing (False Discovery Rate, FDR (Benjamini and Hochberg, 1995)), these results seem to not be driven by a single group, as significant *p*-values are recovered also for the chordate classes studied (apart from fish TpG surplus, supp. table S4). In addition, all arthropod orders are insignificant (Supplementary table S4). Therefore, CpG codons seem to affect codon usage patterns only in chordates, but not in arthropods.

## 3 Discussion

Our study introduces a mechanistic model of codon evolution that can distinguish between selection and mutational biases while accounting for GC–biased gene conversion. Previous research has introduced mechanistic models to estimate mutation and selection coefficients based on population dynamics (e.g., Sharp et al. (2005); Yang and Nielsen (2008); Zeng (2010)) or protein synthesis rates and protein structures (Cope and Gilchrist, 2022; Gilchrist et al., 2015). However, our model, DECUB, also takes into account the effects of gBGC. gBGC has been identified in vertebrates (Figuet et al., 2015) and other insect species (Kent et al., 2012; Wallberg et al., 2015), and there is increasing evidence that its impact is widespread across all metazoans (Pessia et al., 2012; Galtier et al., 2018) and failure to account for it can lead to over or underestimation of selection on codon usage (De Oliveira et al., 2021; Cope and Shah, 2022). DECUB is modelled similarly to the FMutSel model introduced by Yang and Nielsen (2008) and the Moran birth–death process highlighted in Sella and Hirsh (2005). However, in addition to modelling gBGC, our model expands the parameter space to include fitness coefficients of the stop codons, which have not been accounted for in previous studies. This incorporation highlighted a higher fitness of the stop codon TGA in vertebrates, supporting previous studies on the effect of gBGC on the mammalian stop codons (Ho and Hurst, 2022; Trexler et al., 2023).

While DECUB was utilised in this study to investigate phylum–specific patterns of codon usage, it has applications beyond this. It can be implemented on a more refined taxonomic level, such as species families or genera, or even on a per–species basis, enabling us to infer adaptations in codon usage specific to particular lineages. Additionally, DECUB has the potential to investigate sudden changes in the patterns of codon usage, such as those resulting from modifications in the genomic base composition or lineage–specific alterations to the genetic code.

Expanding previous research that studied a small number of representatives per phylum, DECUB was applied here to a vast dataset of over 600 species of chordates and arthropods. We found that codon usage is more extensive in chordates compared to arthropods (fig. 2). Behura and Severson (2012) showed that dipteran and hymenopteran insects have a low extent of codon usage, which is in agreement with our results. Additionally, they also suggested that insects have a pattern of codon usage that is unique to each species, explaining the variation we also observed within arthropods.

Although the extent of codon usage bias varies between chordates and arthropods, we showed that, in both, mutational biases have a more significant impact on shaping genome–wide patterns of codon usage than codon fitnesses (fig. 4). Corroborating these results, the global codon patterns appear to be dominated by a combination of mutational biases towards AT and gBGC also in yeast (LaBella et al., 2019). Mutations have been described previously as the main driver of codon usage in vertebrates (e.g., Doherty and McInerney (2013)), however, their effect on arthropod patterns is less extensive compared to natural selection (Kliman and Hey, 1994).

Our analysis revealed a strong negative correlation between the GC mutational biases – which also include the effects of gBGC – and selection forces acting on the GC–content of each codon (fig. 3). Indeed, despite an excess of GC to AT mutations, GC alleles are more likely to be fixed in mammals due to gBGC or GC–biased selection (Smith and Eyre-Walker, 2001; Behura and Severson, 2013). Birds are known to exhibit similar patterns of gBGC to mammals (Duret and Galtier, 2009; Figuet et al., 2015). However, there is also growing evidence that this phenomenon affects fish (Escobar et al., 2011), whereas its impact on reptiles appears to be less significant (Figuet et al., 2015). These studies support our estimations of mutational biases driven by mutations and gBGC, where reptiles and amphibians exhibited smaller biases towards GC compared to the other chordate classes. In insects, Vicario et al. (2007) found that most genes in *Drosophila* are under GC–biased selection, while Behura and Severson (2012) reported that GC–biases affect codon usage in Diptera, whereas AT–biases are more pronounced in Hymenoptera. This observation corroborates our analysis on arthropods, where mutational biases in Diptera were GC–biased, whereas Hymenoptera was the only taxon with increased AT–bias favouring codons with a higher GC–content.

Finally, our results showed a depletion of CpG and an excess of TpG codons compared to the expected GC–content in chordates but not in arthropods (fig. 5). Methylation in CpG dinucleotides has been shown to hypermutate into TpG through deamination of their cytosine, resulting in a depletion of CpG sites and a surplus of TpG (Simmen, 2008; Miyahara et al., 2015). Most CpG sites are methylated in vertebrates (Bird, 2002), however among arthropods, dipterans exhibit minimal to absent levels of methylation, while other holometabola species (i.e. lepidopterans and hymenopterans) show methylation, but in reduced levels in their protein-coding sequences (Provataris et al., 2018). In contrast, Jabbari and Bernardi (2004) have suggested that differences in GC–content and not methylation may cause CpG shortages. However, our analysis actively compared CpG codons with their expected GC content and found significant differences in chordates, which showcases that methylation in CpG sites rather than GC–content is driving these differences between the two phyla. This difference due to methylation on dinucleotides acting on chordates but not in arthropods can partly explain the more extensive variation in the codon usage patterns in chordates (fig. 2).

In summary, despite differences between chordates and arthropods, in both, mutational biases have a significant impact on shaping genome–wide patterns of codon usage. In all taxa, as GC–mutational biases increase, GC–rich codons become less favourable and vice versa for AT–biased mutations. These contrasting patterns are highlighted in fishes where a strong GC–mutation bias has the most deleterious effect on GC–rich codons, while in some hymenopterans the opposite pattern is observed (fig. 3). This inverse relationship between the mutations towards GC and the fitness coefficients is not merely an artefact of the model (fig. 3), while it also cannot be explained by methylation in arthropods (fig. 5).

A possible explanation might be that stabilising selection is preventing an excess of GC or AT content in the coding regions of the genome, which would limit the occurrence of mRNAs that are either too GC- or AU-rich. Indeed, data from humans, chicken, and *Drosophila* show that GC–content in mRNAs ranges between 30%–70% (Courel et al., 2019). However, we must exercise caution when considering this hypothesis since, apart from methylation in CpG sites, we cannot differentiate selection for translational efficiency in our fitness coefficients from other mutational biases that cannot be captured by our model. Testing this hypothesis may require comparing the mutation rates between GC and AT alleles, gBGC, and codon substitutions rates across the chordate or arthropods phylogeny to determine whether these have co–evolved to stabilise GC content on the coding regions. However, this validation presents challenges and it necessitates polymorphic data as well as experimentally obtained mutation and gBGC rates, which are currently unavailable for most non–model organisms included in our analysis.

This study betters our understanding of the molecular mechanisms involved in the determination of codon composition in animals, the extent of which seems to vary considerably. It also provides insights into the variations in synonymous sites in light of these mechanisms. It is well known that failure to account for this variation breaks the assumption of neutral evolution of synonymous sites and can bias estimates of the ratio of nonsynonymous to synonymous substitution rates, i.e. *ω* = *dN/dS* (Goldman and Yang, 1994; Muse and Gaut, 1994; Spielman and Wilke, 2015), a parameter commonly used to detect natural selection acting on the protein. Therefore, our results highlight the importance for clade–specific approaches in the study of variation at synonymous sites and the detection of natural selection.

## 4 Methods

### 4.1 A model of codon evolution

To assess the evolutionary impact of the forces that govern codon usage bias, we devised a population genetic model on the 64 codons using a Moran model with reversible mutations and selection (supplementary text S1). Mutations are reversible, biased, and modelled as the general time–reversible (GTR) substitution model (Tavaré, 1986), where the mutation rates are proportional to the stationary frequencies *π* between the four nucleotides. GC–biased gene conversion is incorporated as a selection coefficient *γ* favouring GC–alleles (Nagylaki, 1983). Genetic drift is modelled according to the Moran model (Moran, 1958) in a population of *N* individuals, where in each generation one individual is chosen to reproduce and one to die. Finally, selection acting on codons is modelled as a relative fitness coefficient *ϕ*. We further derived the stationary distribution of this model, which defines the frequencies of each of the 64 codons in terms of the aforementioned forces:

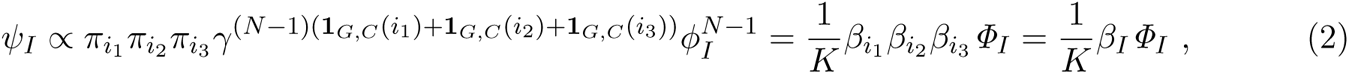

where *i*_1_*i*_2_*i*_3_ are the three nucleotides of codon *I* and **1***_G,C_*(*i*) is the indicator function of nucleotide *i*. The indicator function guarantees that the *γ* parameter contributes to a given codon frequency only if it has a GC nucleotide, thus modelling for gBGC.

Using the stationary distribution of eq. (2), we developed DECUB, a Bayesian estimator that estimates the effects of different evolutionary forces on codon usage from codon counts. As we cannot disentangle them, we combined them into a single parameter *β*, representing mutational biases. Finally, the normalisation factor *K* is the sum of the stationary frequencies of all codons *k*, with 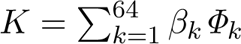such that the stationary distribution adds up to 1.

We normalised each *β* by *π_A_*, the stationary frequency of Adenine, which does not include the effects of gBGC. Similarly, the fitness coefficient *Φ_I_* = *ϕ_I_^N−^*^1^ encompasses the effect of selection with genetic drift and is normalised by *Φ*_ATG_, the fitness coefficient of methionine, as methionine is an essential amino acid and is encoded by a single codon, therefore it is not confounded with codon usage bias. In summary,

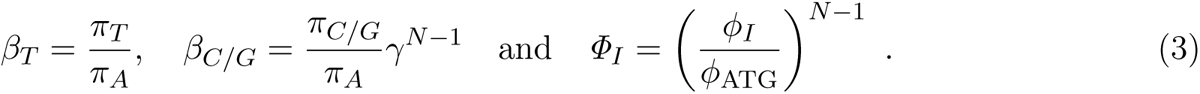

### 4.2 Mapping and coefficient estimation

Mapping is the process of assigning a coefficient to be estimated by the model to a single codon or a group of codons. This can refer to an amino acid mapping, where every coefficient is placed per amino acid and assumes no variation within synonymous codons, or a more codon–usage–specific mapping. We started with an amino acid mapping that includes 23 fitness categories, one for each amino acid and one for each stop codon. Then, for each codon, we calculated the number of species where the predicted frequency is outside the 0.05-0.95 interval of the empirical codon frequency, based on the posterior predictive checks by Gelman and Hill (2006). In a stepwise manner, we added an extra fitness coefficient per amino acid to the codon with the highest error. We continued this process until all codons had an error smaller than 20%, which means that we are predicting its frequency correctly for at least 80% of the species. Using this procedure, we defined a phylum–specific codon mapping for chordates and arthropods, which we incorporated in DECUB, along with the amino acid one. The posterior mean and standard deviation of all parameter estimates of the amino acid and final codon mappings (two chains per mapping) can be found in the supplementary table S1.

The procedure of adding one extra fitness coefficient to each amino acid was repeated in total four times, resulting in four mappings: i.e., the amino acid one, and three codon specific. To compare the fit of these mappings, we employed the Bayesian and Deviance Information Criteria (BIC and DIC) (Schwarz, 1978; Spiegelhalter et al., 2002), where we aimed to find the model with the best fit (lowest score). We then calculated the difference between all the models and the optimal ones (e.g., Δ*DIC* = *DIC_model_ −DIC_optimal_*) and only accepted if the difference was larger than 10 (Spiegelhalter et al., 2002) (supplementary table S2). As chordates reached the identifiability threshold, we could not add more fitness coefficients. For arthropods, we stopped at the last phylum-specific mapping because the Δ*DIC* values between that and the previous one were close to 0, suggesting further mappings would have been redundant (suppl. fig. S10).

We employed DECUB which uses a Bayesian estimator to estimate all mutational and fitness coefficients using the final phylum–specific mapping of chordates and arthropods. We then proceeded to group the estimations of A and T mutational biases together, similarly for G and C, and then group the fitness coefficients by their codon GC–content (fig. S7, S8). Using a linear model, we calculated the slope of fitness coefficients with increasing GC–content. We then calculated the Spearman Rank Correlation Coefficient (Spearman, 1904) between this and the difference between G/C and A/T mutational biases. To test for phylogenetic non-independence which can influence the correlation coefficients and *p*-values, we used the Moran’s I coefficient from the phylosignal package in R (Gittleman and Kot, 1990; Keck et al., 2016). We then calculated phylogenetic contrasts using the ape package in R (Felsenstein, 1985; Paradis and Schliep, 2019) for chordates and arthropods, as well as for each respective order and class based on their respective phylogenies. These contrasts were used to calculate the Spearman’s *ρ* coefficient.

### 4.3 Model validation through simulations

We conducted extensive simulations to establish that our model (1) accurately represents the underlying processes we are modelling and (2) assess our ability to capture genome-wide trends, considering the presence of across genes spatial heterogeneity. We used genome-wide data from humans (*Homo sapiens*) and fruit flies (*Drosophila melanogaster*) as a representative of each of our phyla in our study as there is a plethora of data available for these model organisms. We estimated nucleotide frequencies and combined them with estimates of gBGC to calculate the mutational bias parameters (*β*) for each gene based on equation (3). For humans, we calculated nucleotide frequencies across the majority of human autosomes (after removing outliers) obtained from Ensembl (Accession Number: GCA_000001405.29; Martin et al. (2022)). gBGC was sampled for each gene based on Glémin et al. (2015), where they provided distributions of gBGC for inside and outside hotspots (see Fig. 7 in their publication). We simulated 2% of the genes in hotspots, as typically recombination hotspots are found in 1-2% of the whole genome (Glémin et al., 2015). To calculate the variation in codon fitness, we obtained the whole CDS also from Ensembl (Accession Number: GCA_000001405.29) and derived amino acid preferences relative to methionine, introducing, on top, codon-specific variation based on differences between synonymous codons of the same amino acid. For fruit flies, we calculated nucleotide frequencies across the chromosome arms 2L, 2R, 3L, and 3R obtained from FlyBase (release FB2023_06; Gramates et al. (2022)) and the codon fitnesses based on the CDS from the same release. Estimates of gBGC were obtained from Jackson and Charlesworth (2021). Finally, we utilised these estimates to generate codon counts for 10,000 genes in fruit flies and 20,000 in humans for a total of 100 simulations. The combined codon counts were then input into DECUB to estimate parameters, which were then compared with the gene-wide average simulated values.

### 4.4 Dataset information

Codon counts were collected from the Codon Statistics Database (Subramanian et al., 2022) for a total of 606 species, 415 Chordata and 191 Arthropoda. All species are encoded using the standard genetic code (translation table 1 (Osawa et al., 1992)). The dataset includes counts for all 64 codons, including three stop codons along with the sense ones.

The dataset for chordates contains species from 13 taxa, most of those being shown at the Class level, with most subsequent analyses focusing on a subset of these classes, namely Mammals, Birds, Fish, and all Reptiles and Amphibians combined, which represent almost 97% of all chordates dataset. Similarly, for arthropods, we have 12 taxa of species, the orders of which diverged at similar time points to chordates’ classes. Here, we focused on the orders Diptera (flies and mosquitoes), Lepidoptera (butterflies and moths), and Hymenoptera (ants, bees, and wasps), which represent the majority of species in Hexapoda and comprise 74% of the arthropods dataset (Fig. 6).

### 4.5 Entropy and sensitivity analysis

To measure the evolutionary significance of the mutation and selection bias in the codon frequencies, we calculated the impact of perturbing each of those parameters by *±* 10% on the predicted frequencies. To summarise this effect, we used the Shannon entropy (Shannon, 1948):

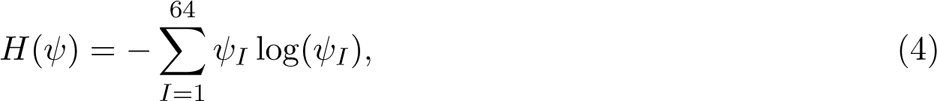

where *ψ_I_* is the predicted frequency of codon *I*. After obtaining the entropy we calculated the relative difference as:

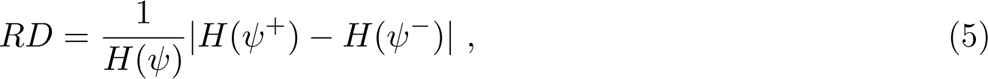

where *ψ*^+^ and *ψ^−^* are the recalculated stationary distribution when we increased and decreased each parameter by 10%, respectively.

### 4.6 Computer statement, Data and Software availability

All analyses were performed in python 3.9.13 (Van Rossum and Drake, 2009), R 4.2.0 (R Core Team, 2022) and C++11 (Josuttis, 2012) with a g++ compiler. DECUB is a free and open–source software written in c++, available on GitHub along with a tutorial on how to use it. The link to the software can be found here: https://github.com/JaneK27/decub.

## Acknowledgements

We thank all members of the Institute of Population Genetics for their feedback and support. This research was funded by the Austrian Science Fund (FWF) [P34524-B] awarded to Rui Borges.

