## Supplementary Materials for "The patterns of codon usage between chordates and arthropods are different but co-evolving with mutational biases"

### Supplementary material

#### Supplementary Text

##### S1. The model

Our model is modelled as a continuous–time Markov process based on Polymorphism–aware Phylogenetic Models, and it includes monomorphic and polymorphic states. Every polymorphic pair is biallelic and differs at a single position, as mutation rates are expected to be much lower than genetic drift in eukaryotes. Mutations are reversible and biased, as the mutation rate from codon $I$ to $J$ ($\mu_{IJ}$) is not necessarily equal to the opposite mutation rate ($\mu_{JI}$). They are assumed to happen only to escape a monomorphic state (boundary sites) and are modelled similarly to the general time–reversible (GTR) substitution model, where the mutation rates are the product of the stationary frequencies ($\pi$) and the exchangeability rates ($\rho$) between the four nucleotides. Genetic drift is incorporated according to the Moran Model and GC–bias as a selection coefficient favouring GC–alleles to capture the effects of GC–biased gene conversion (gBGC). Finally, selection is modelled as a relative fitness coefficient ($\phi$).

If we assume two codons $I = i_1 i_2 i_3$ and $J = j_1 j_2 j_3$ differ at a singe position $c$, with $i_k = j_k \forall k \neq c$, then the transitional matrix $Q$ is:

$$Q = q_{\substack{\{nI,(N-n)J\},\\ \{mI,(N-m)J\}}} = \begin{cases} \mu_{IJ} = \rho_{i_c j_c}\pi_J = \rho_{i_c j_c}\pi_{j_1}\pi_{j_2}\pi_{j_3}, & \text{if } n = N \text{ and } m = N-1 \\ \mu_{JI} = \rho_{j_c I_c}\pi_I = \rho_{j_c I_c}\pi_{i_1}\pi_{i_2}\pi_{i_3}, & \text{if } n = 0 \text{ and } m = 1 \\ \frac{n(N-n)}{N[n\gamma^{\mathbf{1}_{G,C}(i_c)}\phi_I + (N-n)\gamma^{\mathbf{1}_{G,C}(j_c)}\phi_J]}\gamma^{\mathbf{1}_{G,C}(j_c)}\phi_J, & \text{if } 1 \leq n \leq N-1 \text{ and } m = n-1 \\ \frac{n(N-n)}{N[n\gamma^{\mathbf{1}_{G,C}(i_c)}\phi_I + (N-n)\gamma^{\mathbf{1}_{G,C}(j_c)}\phi_J]}\gamma^{\mathbf{1}_{G,C}(i_c)}\phi_I, & \text{if } 1 \leq n \leq N-1 \text{ and } m = n+1 \\ 0, & \text{otherwise} \end{cases} \tag{1}$$

where $n$ and $N-n$ describe the absolute frequencies of the two codons in the initial population of size $N$, $m$ and $N-m$ are the derived frequencies, $\pi_I$ is the nucleotide frequencies for codon $I$ ($\pi_I = \pi_{i_1}\pi_{i_2}\pi_{i_3}$) and $\phi_I$ is the fitness coefficient of codon $I$ (similarly for codon $J$). Finally, $\mathbf{1}_{G,C}(i)$ is the indicator function of nucleotide $i$, which guarantees that the $\gamma$ parameter contributes to a given codon frequency only if it has a GC nucleotide, thus modelling for gBGC.

##### S2. Stationary distribution

The global balance equation characterises the stationary distribution of a Markov chain, and states that the transition rates in and out of state $s_i$ are equal.

$$\psi_{s_i} \sum_{\forall s_i \neq s_j} q_{s_i,s_j} = \sum_{\forall s_i \neq s_j} \psi_{s_j} q_{s_j,s_i} \ , \tag{2}$$

where $\psi_{s_i} q_{s_i,s_j}$ represents the probability flux from state $s_i$ to state $s_j$. So the left–hand side represents the total flow from out of state $s_i$ into states other than $s_i$, while the right–hand side represents the total flow out of all states $s_j \neq s_i$ into state $s_i$.

In a continuous time Markov chain with a transition rate matrix $Q = q_{s_i,s_j}$, where $q_{s_i,s_i} = -q_{s_i}$ and $\psi Q = 0$, the sum of all stationary distributions is equal to 1, $\sum \psi_{s_i} = 1$. When the model is reversible, the global balance equation can be simplified to the detailed balance equations for every pair of states

$s_i$ and $s_j$:

$$\psi_{s_i} q_{s_i,s_j} = \psi_{s_j} q_{s_j,s_i} \tag{3}$$

To calculate the stationary distributions, we will use the rate matrix defined in eq. (1). Assuming we have two codons $I$ and $J$ in a population of $N$ individuals, we have all frequency shifts defined as $q_{\{nI,(N-n)J\},\{mI,(N-m)J\}}$. Initially, as a first step assuming we start from a population of fixed for the codon $I$ individuals ($\{NI\}$), going from the monomorphic state to the first polymorphic state ($\{(N-1)I, J\}$), based on eq. (3), we get:

$$\begin{aligned} &\psi_{\{NI\}} q_{\{NI\},\{(N-1)I,J\}} = \psi_{\{(N-1)I,J\}} q_{\{(N-1)I,J\},\{NI\}} \\ &\Leftrightarrow \psi_{\{NI\}} N \rho_{i_c j_c} \pi_J = \psi_{\{(N-1)I,J\}} \frac{(N-1)}{N[(N-1)\gamma^{\mathbf{1}_{G,C}(i_c)}\phi_I + \gamma^{\mathbf{1}_{G,C}(j_c)}\phi_J]} \gamma^{\mathbf{1}_{G,C}(i_c)} \phi_I \end{aligned} \tag{4}$$

Similarly, the same will hold going from $\{(N-1)I, J\}$ to $\{(N-2)I, 2J\}$ and opposite.

$$\begin{aligned} &\psi_{\{(N-1)I,J\}} q_{\{(N-1)I,J\},\{(N-2)I,2J\}} = \psi_{\{(N-2)I,2J\}} q_{\{(N-2)I,2J\},\{(N-1)I,J\}} \\ &\Leftrightarrow \psi_{\{(N-1)I,J\}} \frac{(N-1)}{N[(N-1)\gamma^{\mathbf{1}_{G,C}(i_c)}\phi_I + \gamma^{\mathbf{1}_{G,C}(j_c)}\phi_J]} \gamma^{\mathbf{1}_{G,C}(j_c)} \phi_J = \\ &\psi_{\{(N-2)I,2J\}} \frac{2(N-2)}{N[(N-2)\gamma^{\mathbf{1}_{G,C}(i_c)}\phi_I + 2\gamma^{\mathbf{1}_{G,C}(j_c)}\phi_J]} \gamma^{\mathbf{1}_{G,C}(i_c)} \phi_I \\ &\Leftrightarrow \psi_{\{(N-1)I,J\}} \frac{(N-1)}{N[(N-1)\gamma^{\mathbf{1}_{G,C}(i_c)}\phi_I + \gamma^{\mathbf{1}_{G,C}(j_c)}\phi_J]} = \\ &\psi_{\{(N-2)I,2J\}} \frac{2(N-2)}{N[(N-2)\gamma^{\mathbf{1}_{G,C}(i_c)}\phi_I + 2\gamma^{\mathbf{1}_{G,C}(j_c)}\phi_J]} \frac{\gamma^{\mathbf{1}_{G,C}(i_c)}}{\gamma^{\mathbf{1}_{G,C}(j_c)}} \frac{\phi_I}{\phi_J} \end{aligned} \tag{5}$$

Combining the equations (5) and (4), we get:

$$\begin{aligned} &\psi_{\{NI\}} N \rho_{i_c j_c} \pi_J = \psi_{\{(N-2)I,2J\}} \frac{2(N-2)}{N[(N-2)\gamma^{\mathbf{1}_{G,C}(i_c)}\phi_I + 2\gamma^{\mathbf{1}_{G,C}(j_c)}\phi_J]} \frac{\gamma^{\mathbf{1}_{G,C}(i_c)}}{\gamma^{\mathbf{1}_{G,C}(j_c)}} \frac{\phi_I}{\phi_J} \gamma^{\mathbf{1}_{G,C}(i_c)} \phi_I \\ &\Leftrightarrow \psi_{\{NI\}} N \rho_{i_c j_c} \pi_J = \psi_{\{(N-2)I,2J\}} q_{\{(N-2)I,2J\},\{(N-1)I,J\}} \frac{\gamma^{\mathbf{1}_{G,C}(i_c)}}{\gamma^{\mathbf{1}_{G,C}(j_c)}} \frac{\phi_I}{\phi_J} \\ &\Leftrightarrow \psi_{\{NI\}} N \rho_{i_c j_c} \pi_J \frac{\gamma^{\mathbf{1}_{G,C}(j_c)}}{\gamma^{\mathbf{1}_{G,C}(i_c)}} \frac{\phi_J}{\phi_I} = \psi_{\{(N-2)I,2J\}} q_{\{(N-2)I,2J\},\{(N-1)I,J\}} \end{aligned} \tag{6}$$

Generalising eq. (6) for state $\{(N-n)I, nJ\}$, we get:

$$\psi_{\{NI\}} N \rho_{i_c j_c} \pi_J \frac{\gamma^{\mathbf{1}_{G,C}(j_c)(n-1)}}{\gamma^{\mathbf{1}_{G,C}(i_c)(n-1)}} \frac{\phi_J^{(n-1)}}{\phi_I^{(n-1)}} = \psi_{\{(N-n)I,nJ\}} q_{\{(N-n)I,nJ\},\{(N-n-1)I,(n+1)J\}} \tag{7}$$

Finally, taking it all the way to the last state ($\{NJ\}$), we have:

$$\psi_{\{NI\}} N \rho_{i_c j_c} \pi_J \frac{\gamma^{\mathbf{1}_{G,C}(j_c)(N-1)}}{\gamma^{\mathbf{1}_{G,C}(i_c)(N-1)}} \frac{\phi_J^{(N-1)}}{\phi_I^{(N-1)}} = \psi_{\{NJ\}} q_{\{NJ\},\{I,(N-1)J\}} \Leftrightarrow \frac{\psi_{\{NI\}}}{\psi_{\{NJ\}}} = \frac{\pi_I}{\pi_J} \frac{\gamma^{\mathbf{1}_{G,C}(i_c)(N-1)}}{\gamma^{\mathbf{1}_{G,C}(j_c)(N-1)}} \frac{\phi_I^{N-1}}{\phi_J^{N-1}} \tag{8}$$

To generalise for all monomorphic states, we need to include all codon positions in the GC–bias parameter. Since we are only studying the frequency shifts between codons that differ in a single position, the other two codon positions are the same, therefore have equal GC–biases. In equation (8) the $\gamma$ parameter can be expanded to:

$$\frac{\gamma^{\mathbf{1}_{G,C}(i_c)(N-1)}}{\gamma^{\mathbf{1}_{G,C}(j_c)(N-1)}} = \frac{\gamma^{\mathbf{1}_{G,C}(i_1)(N-1)}\gamma^{\mathbf{1}_{G,C}(i_2)(N-1)}\gamma^{\mathbf{1}_{G,C}(i_3)(N-1)}}{\gamma^{\mathbf{1}_{G,C}(j_1)(N-1)}\gamma^{\mathbf{1}_{G,C}(j_2)(N-1)}\gamma^{\mathbf{1}_{G,C}(j_3)(N-1)}} = \frac{\gamma^{(N-1)(\mathbf{1}_{G,C}(i_1)+\mathbf{1}_{G,C}(i_2)+\mathbf{1}_{G,C}(i_3))}}{\gamma^{(N-1)(\mathbf{1}_{G,C}(j_1)+\mathbf{1}_{G,C}(j_2)+\mathbf{1}_{G,C}(j_3))}} \quad (9)$$

By incorporating the generalisation for gBGC in equation (9) with the fraction we got in equation (8), we get the approximation for the stationary frequencies for the monomorphic states.

$$\begin{aligned} \psi_{\{NI\}} &\propto \pi_I \gamma^{(N-1)(\mathbf{1}_{G,C}(i_1)+\mathbf{1}_{G,C}(i_2)+\mathbf{1}_{G,C}(i_3))}\phi_I^{N-1} \\ \psi_{\{NJ\}} &\propto \pi_J \gamma^{(N-1)(\mathbf{1}_{G,C}(j_1)+\mathbf{1}_{G,C}(j_2)+\mathbf{1}_{G,C}(j_3))}\phi_J^{N-1} \end{aligned} \quad (10)$$

#### Supplementary Tables

**Table S3: Species information.** Number of overall species as well as per group tested (top table), and number of species, average number of genes and sites for chordates and artrhopods (bottom table).

| **Phylum** | **Taxon** | | **# of species** |
|---|---|---|---|
| **Chordata** | Total | | 415 |
| | Mammalia | Mammals | 163 |
| | Aves | Birds | 88 |
| | Reptilia & Amphibia | Reptiles & Amphibians | 30 |
| | Actinopteri | Fish | 117 |
| **Arthropoda** | Total | | 191 |
| | Diptera | Flies & Mosquitoes | 64 |
| | Lepidoptera | Butterflies & Moths | 23 |
| | Hymenoptera | Ants, Bees & Wasps | 55 |

| **Phylum** | **# of species** | **Average # of genes** | **Average # of sites** |
|---|---|---|---|
| **Chordata** | 415 | 20,837.3 | 11,792,024 |
| **Arthropoda** | 191 | 14,171.1 | 7,491,224 |

*Note: Reptilia includes the classes including Crocodylia, Testudines, and Lepidosauria.*

**Table S4: Statistical measurements between CpG (and TpG) sites and their respective GC-content.** Values for the one-sample Wilcoxon signed-rank test Statistic and p-value is shown after FDR correction for the groups analysed.

| **Chordata** | | | | | |
|---|---|---|---|---|---|
| | | **CpG <3 GCs** | | **TpG >1 GC** | |
| *Taxon* | | *Statistic* | *p-value* | *Statistic* | *p-value* |
| Overall | | 538226 | <2.2e-16* | 4709110 | <2.2e-16* |
| Mammalia | Mammals | 1050 | <2.2e-16* | 792163 | 1.512e-14* |
| Aves | Birds | 52 | <2.2e-16* | 239470 | 2.12e-12* |
| Amphibia+Reptilia | Amphibians+Reptiles | 33 | <2.2e-16* | 37049 | 1.373e-07* |
| Actinopteri | Fish | 53935 | <2.2e-16* | 355859 | 0.08656 |

| **Arthropoda** | | | | | |
|---|---|---|---|---|---|
| | | **CpG <3 GCs** | | **TpG >1 GC** | |
| *Taxon* | | *Statistic* | *p-value* | *Statistic* | *p-value* |
| Overall | | 25552 | 1 | 139948 | 0.9997 |
| Diptera | Flies+Mosquitos | 6065 | 1 | 23892 | 1 |
| Lepidoptera | Butterflied+Moths | 675 | 1 | 3725 | 1 |
| Hymenoptera | Ants+Bees+Wasps | 4152 | 1 | 23161 | 1 |

#### Supplementary Figures

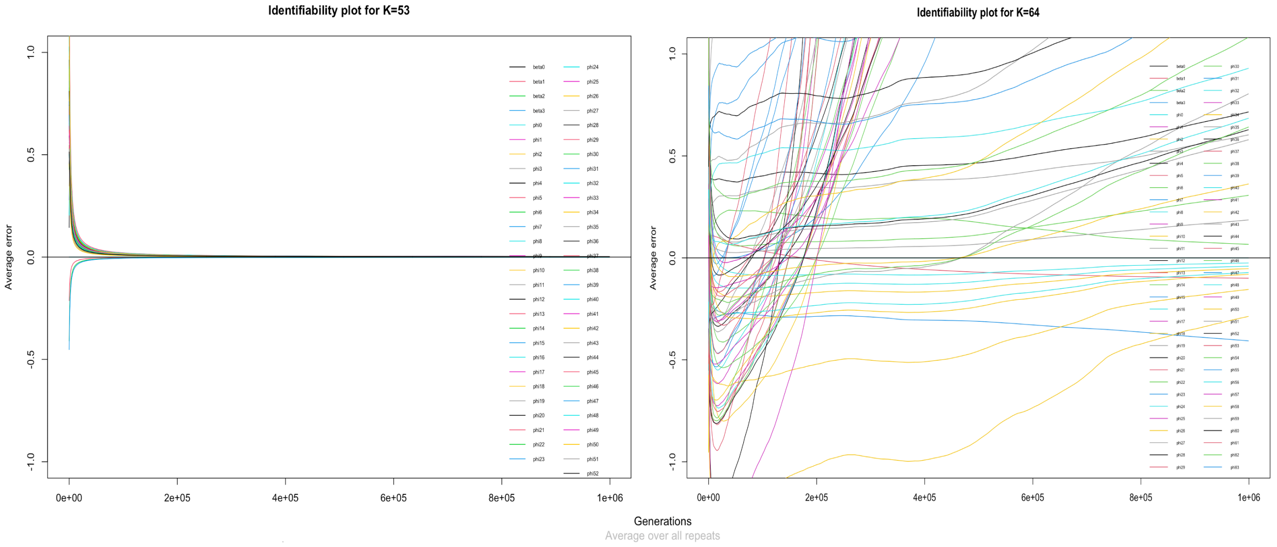


**Figure S1: Cumulative sum of errors of each parameter estimate over the MCMC generations.** The plot on the left represents the cumulative errors of simulated mutational bias parameters and 53 simulated fitness coefficients ($K = 53$; identifiability threshold). After 53 coefficients the model becomes unidentifiable, like the plot on the right with $K = 64$ fitness coefficients (one codon coefficient per codon). Both plots show the cumulative errors averaged over 8 repeats for 1 million MCMC generations.

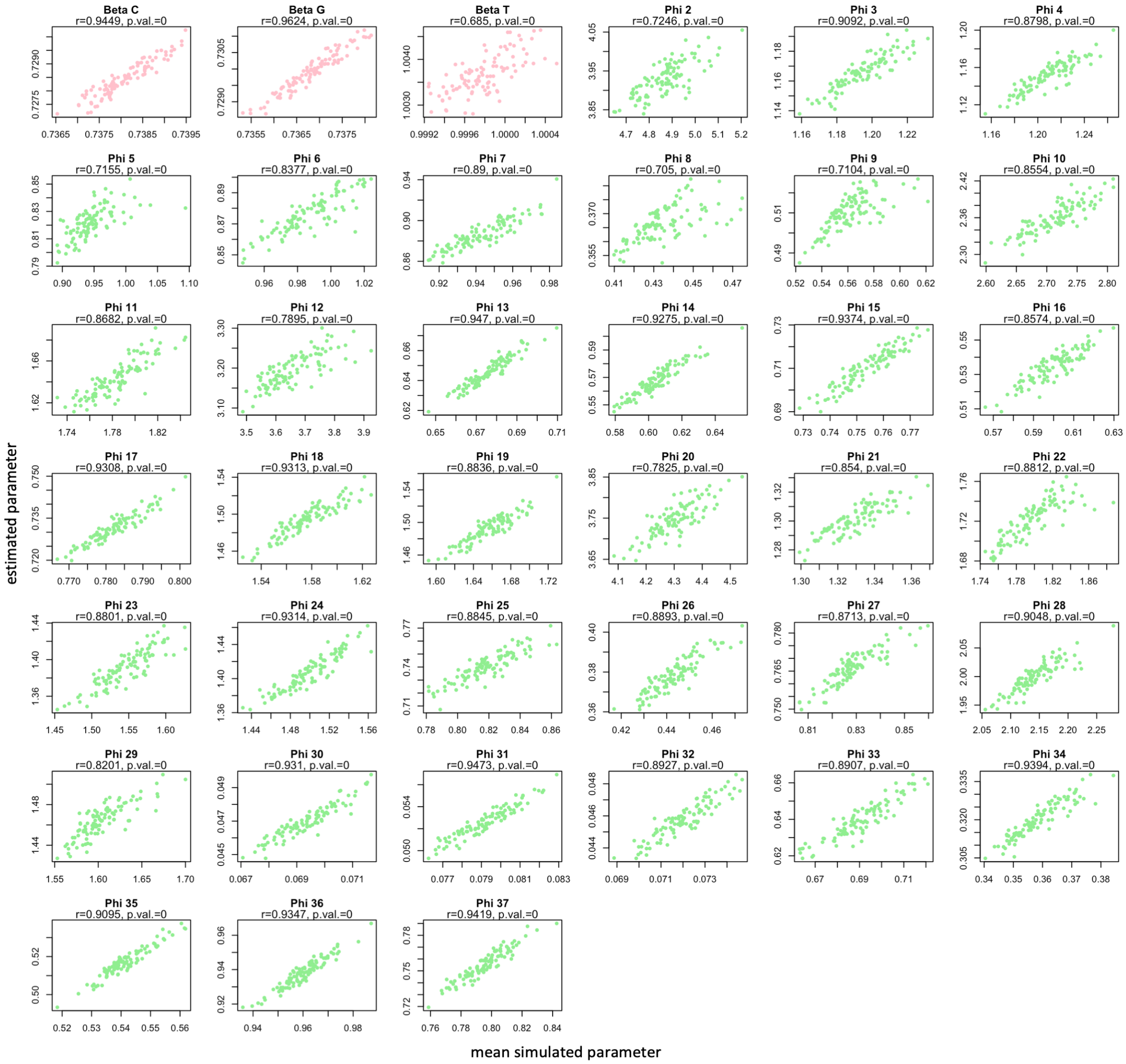


**Figure S2: Relationship between mean simulated values and estimations in *Drosophila melanogaster*.** Comparison of average parameter values simulated across 10,000 fruit fly genes and those estimated by DECUB. Spearman's $\rho$ correlation coefficient is provided alongside its corresponding p-value. Mutational biases are denoted by pink dots, and codon fitnesses are represented by green dots.

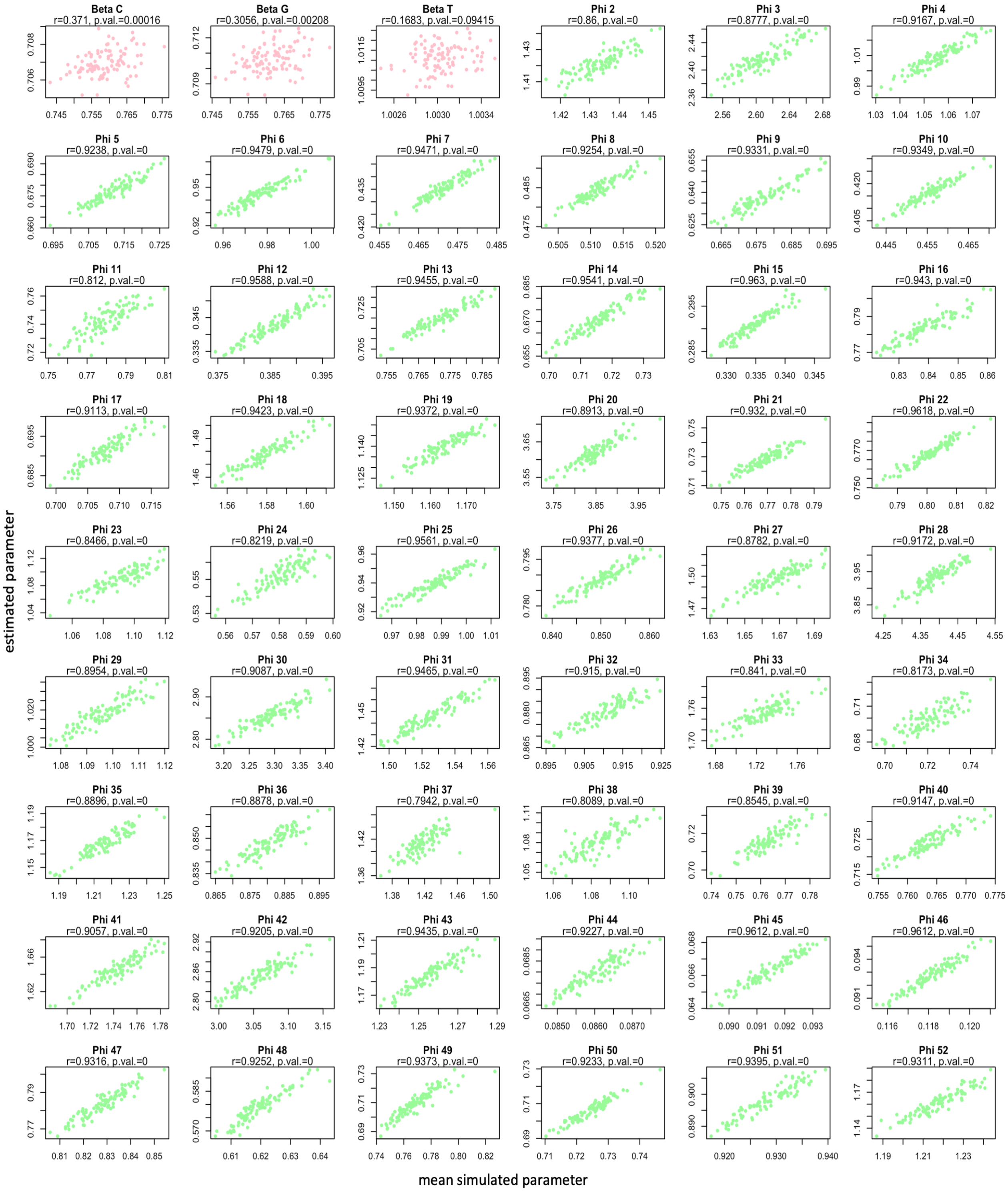


**Figure S3: Relationship between mean simulated values and estimations in *Homo sapiens*.** Comparison of average parameter values simulated across 20,000 human genes and those estimated by DECUB. Spearman's $\rho$ correlation coefficient is provided alongside its corresponding p-value. Mutational biases are denoted by pink dots, and codon fitnesses are represented by green dots.

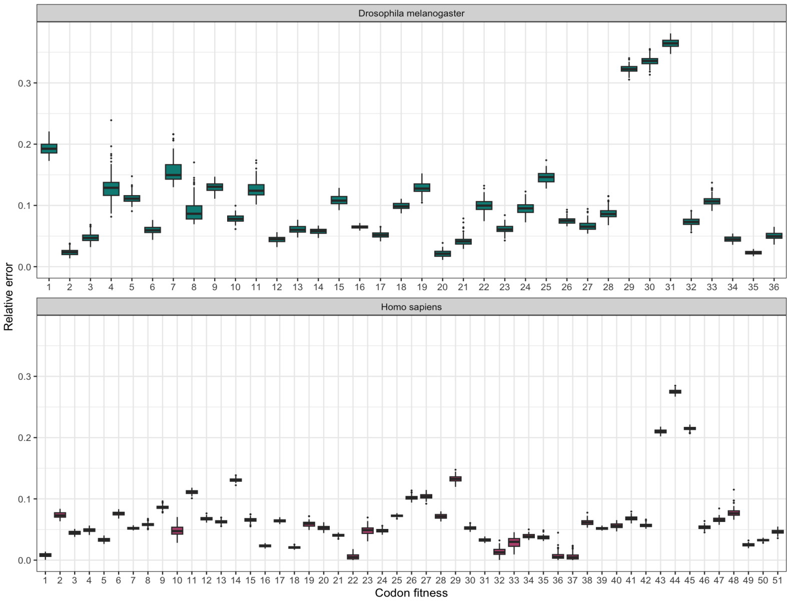


**Figure S4: Relative error between mean simulated codon fitnesses and estimated ones in *Drosophila melanogaster* and *Homo sapiens*.** Relative error between estimated and average simulated codon fitnesses across 10,000 genes in fruit flies and 20,000 genes in humans.

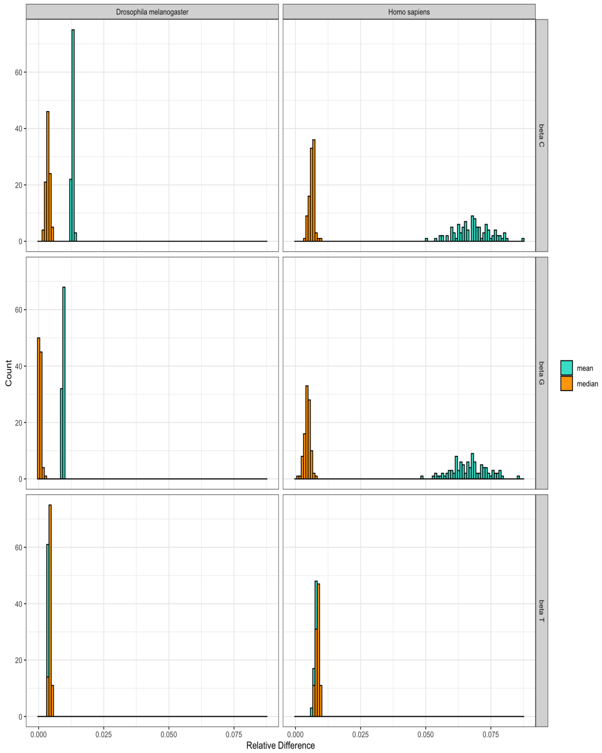


**Figure S5: Comparison of relative errors between mean and median simulated mutational biases and their respective estimated values in *Drosophila melanogaster* and *Homo sapiens*.** Relative error between estimated and mean simulated mutational biases (shown in blue) and the median (shown in orange) across 10,000 genes in fruit flies and 20,000 genes in humans.

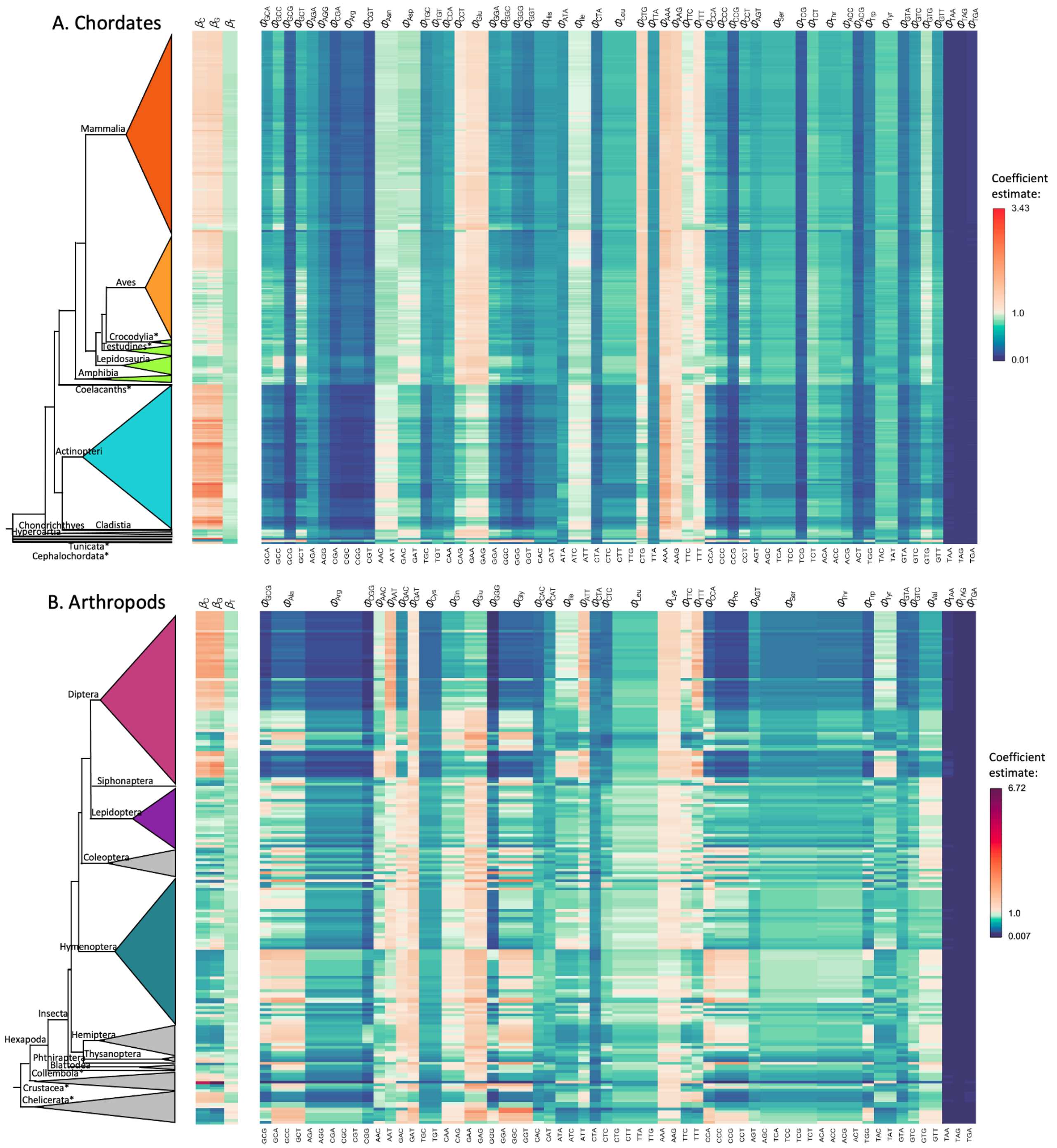


**Figure S6: Image plots of parameter estimates after the final mapping for chordates and arthropods.** Parameter estimate for each mutational bias coefficient $\beta$ and fitness coefficient $\Phi$ (top of image plots). The codons corresponding to the fitness coefficients are shown at the bottom of the image plots and the colours represent the magnitude of the estimate with white being equal to the reference, blue disfavoured and red shows the favoured estimates. Finally, on the left, the corresponding phylogeny of the species is shown for (A) chordates and (B) arthropods.

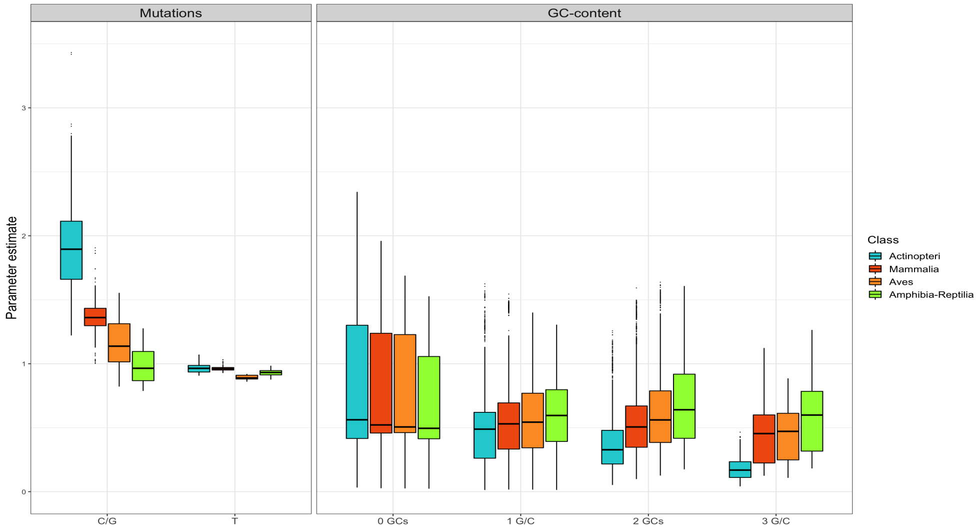


**Figure S7: Mutational biases and fitness coefficient estimates in chordates.** Grouped mutational estimates for C/G and A/T mutational biases and fitness coefficient estimates grouped according to the GC–content of each respective codon. Only the codons with a specific fitness coefficient were considered (excluding fitness coefficients of grouped codons encoding the same amino acid).

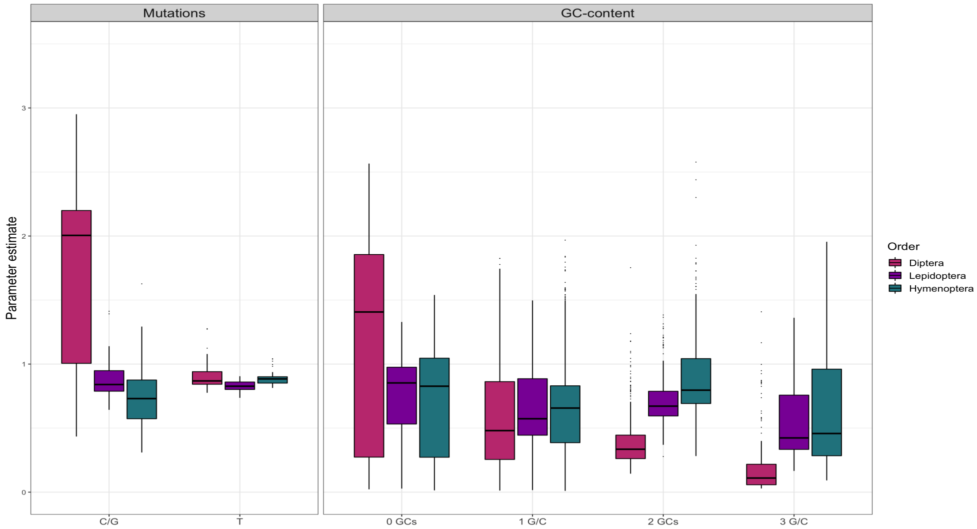


**Figure S8: Mutational biases and fitness coefficient estimates in arthropods.** Grouped mutational estimates for C/G and A/T mutational biases and fitness coefficient estimates grouped according to the GC–content of each respective codon. Only the codons with a specific fitness coefficient were considered (excluding fitness coefficients of grouped codons encoding the same amino acid).

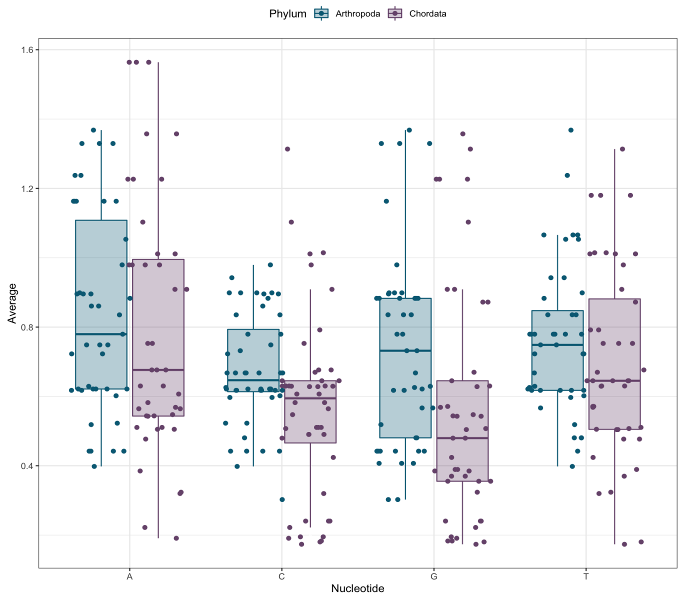


**Figure S9: Distribution of fitness coefficients per nucleotide in arthropods and chordates.** Boxplots showing no association between the distribution of fitness coefficients and their nucleotide composition. This plot helps demonstrate that fitness coefficients are not compensating for the mutation parameters. This comparison was validated using a Wilcoxon test with FDR correction. From the 12 comparisons, only the distribution of fitness coefficients of nucleotides A and G in chordates show significant differences ($p$–value=0.0056).

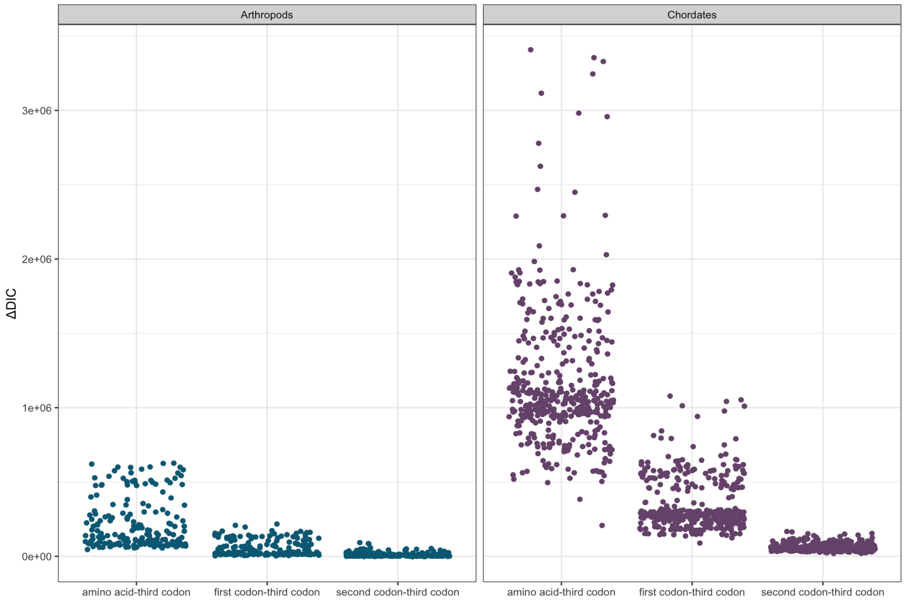


**Figure S10: $\Delta DIC$ values between previous mapping and the last codon mapping.** Comparisons between the previous mappings with the optimal one (last codon mapping, smallest DIC value). In both arthropods and chordates, the last one was preferred with $\Delta DIC$ values $> 10$, the threshold for accepting the optimal model.
